# Modeling science trustworthiness under publish or perish pressure

**DOI:** 10.1101/139063

**Authors:** David Robert Grimes, Chris T. Bauch, John P.A. Ioannidis

## Abstract

The scientific endeavor pivots on the accurate reporting of experimental and theoretical findings, and consequently scientific publication is immensely important. As the number of active scientists continues to increase, there is concern that rewarding scientists chiefly on publication creates a perverse incentive where careless and fraudulent research can thrive. This is compounded by the predisposition of top-tier journals towards novel or positive findings rather than negative results or investigations that merely confirm a null hypothesis, despite their intrinsic value, potentially compounding a reproducibility crisis in several fields. This is a serious problem for both science and public trust in scientific findings. To date, there has been comparatively little mathematical modeling on the factors that influence science trustworthiness, despite the importance of quantifying the problem. In this work, we present a simple phenomenological model with cohorts of diligent, careless and unethical scientists with funding allocated based on published outputs. The results of this analysis suggest that trustworthiness of published science in a given field is strongly influenced by the false positive rate and the pressures from journals for positive results, and that decreasing available funding has negative consequences for the resulting trustworthiness. We also examine strategies to combat propagation of irreproducible science, including increasing fraud detection and awarding diligence, discussing the implications of these findings.

## Introduction

In academia, the phrase “publish or perish” is more than a pithy witticism - it reflects the reality that researchers are under immense pressure to continuously produce outputs, with career advancement dependent upon them [1, 2]. Academic publications are deemed a proxy for scientific productivity and ability, and with an increasing number of scientists competing for funding, the previous decades have seen an explosion in the rate of scientific publishing [3]. Yet whilst output has increased dramatically, increasing publication volume does not imply that the average trustworthiness of publications has improved. A previous paper by Ioannidis [4] has outlined the reasons why many published research findings are false, and the dubious use of P-values for significance in research findings has of late been widely discussed [5–9]. Across much of experimental science from psychology [10] to biomedical science [11–13] and cancer research [14], there is concern over an apparent reproducibility crisis.

Despite their vital importance in conveying accurate science, top-tier journals possess a limited number of publication slots and are thus overwhelmingly weighed towards publishing only novel or significant results. Despite the fact that null results and replications are important scientific contributions, the reality is that journals do not much care for these findings. Researchers are not rewarded for submitting these findings nor for correcting the scientific record, as high profile examples attest [15, 16]. This pressure to produce positive results may function as a perverse incentive. Edwards and Roy [17] argue that such incentives encourage a cascade of questionable findings and false positives. Heightened pressure on academics has created an environment where “*Work must be rushed out to minimize the danger of being scooped”* [18]. The range of questionable behavior itself is wide [19]. Classic ‘fraud’ (falsification, fabrication, and plagiarism (FFP) [20]) may be far less important than more subtle questionable research practices.

So how common are such practices? A study of National Institute of Health (NIH) funded early and mid-career scientists (n = 3247) found that within the previous three years, 0.3% admitted to falsification of data, 6% to a failure to present conflicting evidence and a worrying 15.5% to changing of study design, methodology or results in response to funder pressure [21]. An overview by Fanelli [22] has shown that questionable research practices are as common as 75%, while fraud per se occurs only in 1-3% of scientists. These findings are alarming, yet quantification of these perverse incentives is vital if we are to understanding the potential extent of the underlying problem, and formulate strategies to address it. This is an underdeveloped area, but one which is slowly growing - recent works by Smaldino and McElreath [23, 24] have employed elegant dynamic models to demonstrate that even when there is no attempts at fraud or untoward research practices, selection based solely on published output tends to produce poorer methods and higher false discovery rates, a phenomenon they term “*the natural selection of bad science”*.

Suboptimal science and fraud can take myriad forms which renders it difficult to detect [25]. For the purposes of this article, we define fraud as an explicit ‘intention to deceive’ [26].A more recent investigation [22] put the weighed mean percentage of scientists committing research fraud as high as 1.97%, with over a third admitting to questionable research practices. The same investigation found that about 14.12% of scientists reported observing fraudulent research behavior in colleagues. Another study [27] found that 5% of responding authors claimed to be personally aware of fabricated or misrepresented data in a trial they had participated in. A study of bio-statisticians [28] found that over half of respondents reported being aware of research misconduct.

A 2012 [29] analysis found that FFP offenses rather than honest error accounted for 67.4% of retracted publications, with the rate of retraction due to fraud increasing ten-fold since 1975. An important question is whether scientists who are unethical (fraudulent) or sloppy (careless) may thrive and even outperform diligent scientists in a system driven by publish or perish pressure. Since it is impossible to identify all unethical and careless scientists, one can perform mathematical modeling of science under different assumptions and find out how these scientists fare and what the implications are for the overall trustworthiness of science.

To better understand the impact of publish or perish on scientific research, and to garner insight into what drives the trustworthiness of published science is of paramount importance if we are to counter-act any detrimental impacts of such practices. In this work, we present a simple but instructive model of scientific publishing trustworthiness under the assumption that researchers are rewarded for their published output, taking account of field-specific differences and the proportion of resources allocated with funding cycle. The factors that influence resultant trustworthiness are quantified and discussed, as well as implications for improving the trustworthiness of scientific publishing.

## Model Outline

### Basic model and assumptions

To construct a simple model of publication rewards, we define the total amount of available funding for research as *R*(*t*). Per unit of funding in a given field, there is a global discovery rate of *D_R_*, which includes a proportion *p_T_* of positive / significant results, a proportion *p_F_* of false positives, and a proportion *n* of null results. Null results in principle can include both true negatives and false negatives, but given the bias toward positive results we will not discriminate between these two in this investigation. The relative proportion of positives and nulls will be inherently field-specific - certain disciplines will be more prone to false positives, whilst others tend to yield less ambiguous results. Since the quantities are proportions we have that

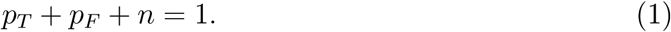

In certain fields, the false positive rate may be high, and so diligent researchers take measures to falsify positive results and test their results multiple times. Even when research groups are very diligent, they may reasonably submit a fraction *ε* of their false positives. Researchers exist on a spectrum, but for simplicity we may broadly sub-divide this spectrum into three distinct classes.

1. Diligent cohort - This group take pains to replicate experiments and do not dishonestly manipulate results. Their false positive submission fraction is *ε*, thus as low as reasonably possible. They account for a fraction *f_D_* of the initial total, and parameters relating to them have subscript *D*.
2. Careless cohort - This group do not falsify results, but are much less careful at eliminating spurious positive results. They may also have questionable practices that lead them to false positives. As a result, they have a false positive submission rate of *cε* where *c* > 1. They account for a fraction *f_C_* of the initial total, and parameters relating to them have subscript *C*.
3. Unethical cohort - This group appear broadly similar to the diligent group, but with one crucial difference in that may occasionally manipulate data or knowingly submit dubious results at a rate of *δ* beyond global discovery rate. For convenience, instead of defining a higher value of *D_R_* in this group to account for the higher “discovery” rate, we retain the same parameter value of *D_R_* for the unethical cohort but allow *p_T_* + *p_F_* + *n* + *δ* > 1, so that their realized “discovery” rate is higher than the other groups. They account for a fraction *f_U_* of the initial total, and parameters relating to them have subscript *U*.

The funding held by the diligent cohort at a given time is *x*(*t*), with *y*(*t*) help by the careless cohort and *z*(*t*) by the unethical cohort, so that

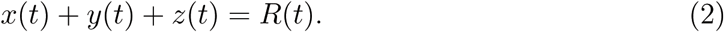

With these assumptions, we can model the theoretical impact of a paradigm where researchers are rewarded with funding and success in direct relation to their publication output. As outlined in the introduction, there is huge pressure on scientists to submit positive or ‘novel’ findings, whilst findings confirming the null hypothesis are frequently side-lined. Under such a selection pressure, all researchers will aim to submit their significant positive results for publication. The respective rates of submission per unit funding for the diligent, careless and unethical cohorts are accordingly

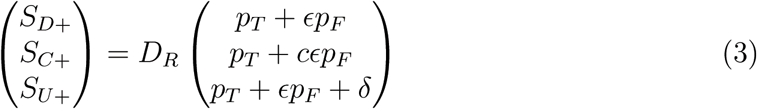

The rate at which null results are submitted is less clear - in general, there is a significant bias in publication towards significant results. As a consequence, negative findings are often shunned by high impact journals, and scientists are disinclined to submit them, meaning that potentially important null results may not ever see the light of publication, the so-called ‘file drawer’ problem. We assume that each cohort submit only a fraction of their null results in the proportions *β_D_*, *β_C_*, *β_UD_* such that

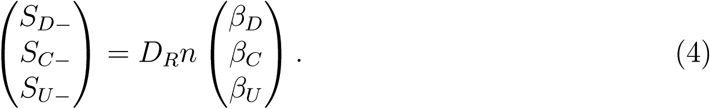

Equations 1–4 comprise the researcher specific parameters, and we must further quantify the journal specific elements also. Competition for space in field-specific top-tier journals is fierce, and we denote the combined carrying-capacity of these field-specific top-tier journals as *J* (*t*). These journals exhibit a clear bias towards positive results, with a positive-publication weighing fraction of published articles, *B*, describing significant results. Thus, we can quantify the probability that a positive result (*ν_P_* (*t*)) or a negative result (*ν_N_* (*t*)) is published. These probabilities are given by

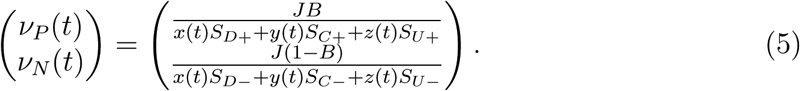

From this, we can then yield an expression for the publication rate per unit of funding for the diligent, careless and unethical cohorts, which are respectively

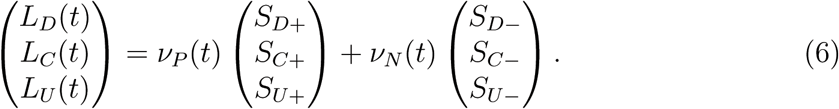

The average rate of publications per unit of funding per unit time is thus

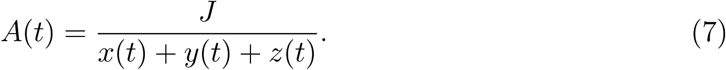

If researchers are rewarded with funding based solely on their published output, we can quantify the impact of this with time by employing a recursive series solution at discrete time-steps, corresponding to funding cycles. If funding is allocated to each cohort is based upon their output at the beginning of the previous funding cycling, and we assume total funding remains constant 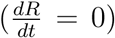 then the funding available for each cohort at each successive time step is

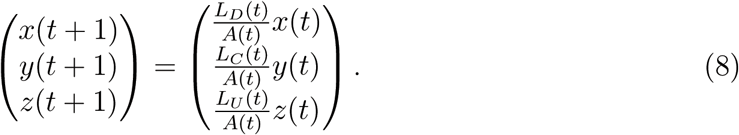

### Variable funding resources

We also consider the fact that the total amount of funding may not remain constant, so we may model the impact of changing funding scenarios. For simplicity, we assume it changes at some constant rate *G*, which can be negative (for diminishing funding, the likes of which might occur with a decrease in NIH or EU funding budgets), zero (for constant funding, as in equation 8) or positive (increasing funding). New funding is allocated at random in proportions reflecting the typical make-up of new researchers, and accordingly the refined equations are

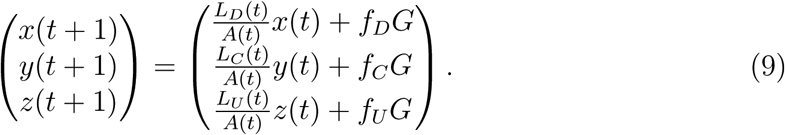

### Research fraud detection

For unethical researchers, we can look at a slightly more complicated scenario where dubious publications have a probability of detection leading to denial of funding, *η*. We further assume this penalization only applies to dubious results which were published rather than just submitted. If this consideration is taken into account, then we modify the last part of equation 9 to reflect this so that

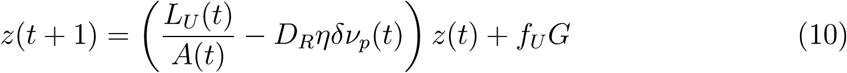

### Rewarding diligence

The diligent cohort have intrinsically lower submission rates than other groups, and consequently are more likely to suffer under a publish or perish regime, despite the importance of their reproducible work. To counter-act this, it has been suggested that rewarding diligence might counteract this trend [30, 31]. We might envision a situation where scientific works are audited for reproducibility, with groups who keep their reproducibility high and error rates below a certain unavoidable threshold (given by *D_R_ν_P_ ε*) garnering a reward of *R_W_*. This in practice could only be achieved by the diligent cohort, and in the most simple case, their funding resources are given by

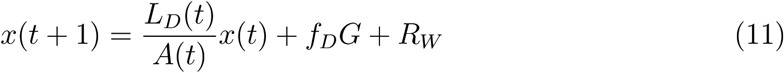

### Counter-acting publication bias

It is also possible to envision a situation where journals don't give any preference to positive results over null results. In this case, we would expect researchers to submit all their results so that *β_D_* = *β_C_* = *β_U_* = 1. In this case, *ν_P_* and *ν_N_* are replaced by a single function of time *ν*, given by

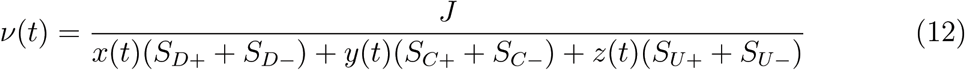

### Trustworthiness of published science

Finally, we define a metric for the trustworthiness of published science, defined as the proportion of reproducible results, *T* (*t*). This is given by

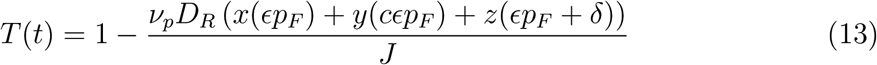

where the time arguments of *x*,*y*,*z* and *ν_p_* have been excluded for clarity.

### Parameter estimation and assumptions

Details on the parameter estimation and assumptions and the range of values considered for each parameter in the model appear in supplement 1 and supplementary table 1.

## Results

### Impact of the field-specific false positive rate

Figure 1 shows the change in funding proportions with time for each cohort in a field with a low rate of false positives 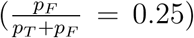 and a field with a high rate of false positives 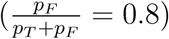. What is immediately evident is that in that in fields where false positives comprise the bulk of positive results, the trustworthiness of published science suffers markedly, and careless and unethical cohorts are disproportionately rewarded at the expense of diligent researchers. Simulation results suggests that the trustworthiness of published science in any given field is strongly dependent on the false positive rate in that field under a publish or perish paradigm.

**Figure 1:**
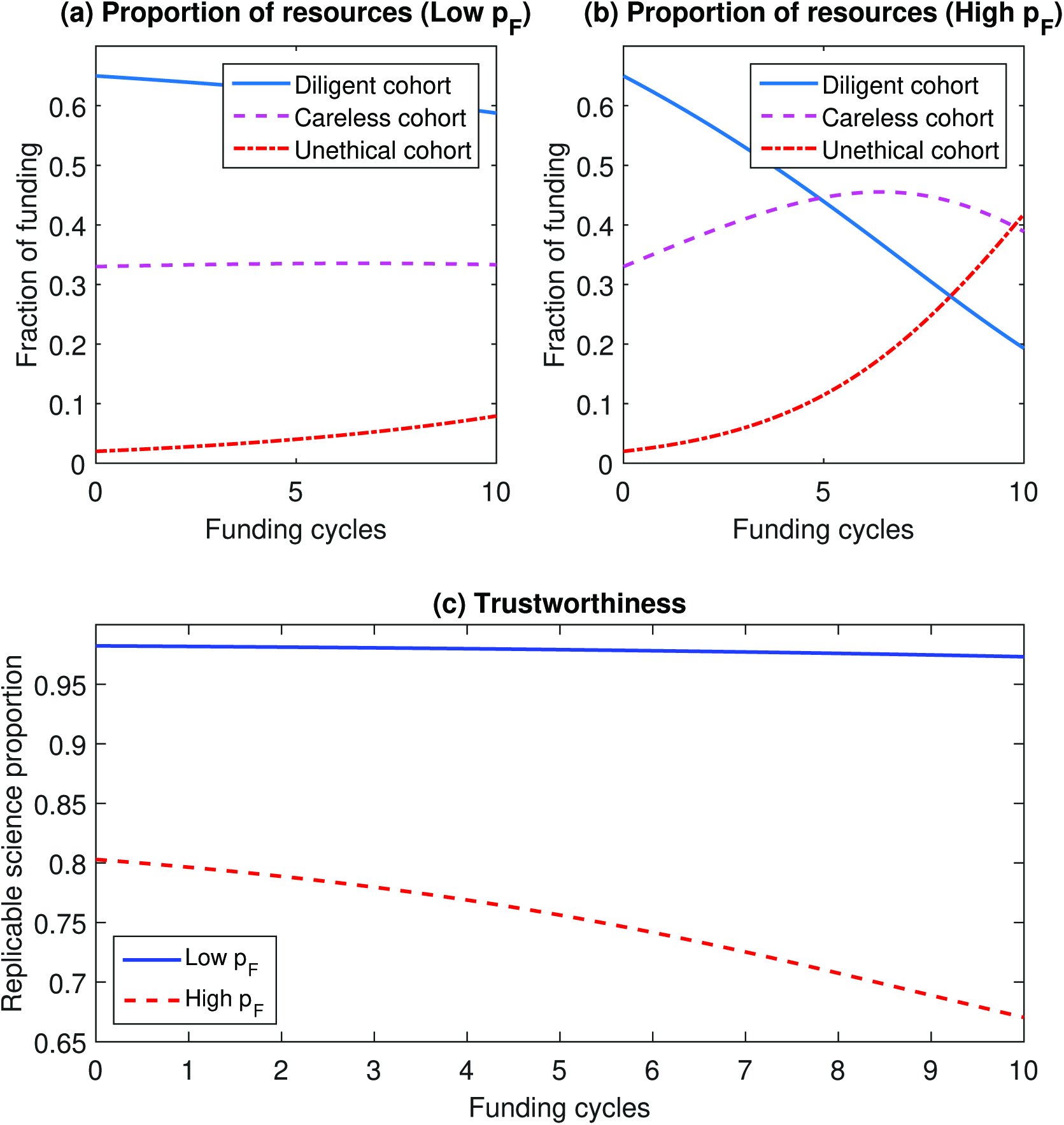
The impact of field-specific false positive rate on resources allocated and science trustworthiness. (a) depicts the projected funding allocations in a field where false positives a re relatively rare (*P_T_* = 0.32,*p_F_* = 0.08). By contr ast, (b) s hows the impact on resources consumed when false positives are the no rm (*P_T_* = 0.08,*p_F_* = 0.32). In (c), the trustworthiness (proportion of repr od uci ble science) for both scenar ios are depicted, indicat ing false positive rate highly drives the trustworthiness of scientific publication in a given field.

### Impact of funding growth rate

As depicted in figure 2. The increasing of available funds has the net effect of reducing publication pressure by bring down the average number of publications expected per unit funding, provided journal capacity stays roughly constant, reducing the likelihood of dubious publications being selected. Conversely, reducing funding increases the publication pressure and results in increases selection of suspect works and a fall in scientific reproducibility.

**Figure 2:**
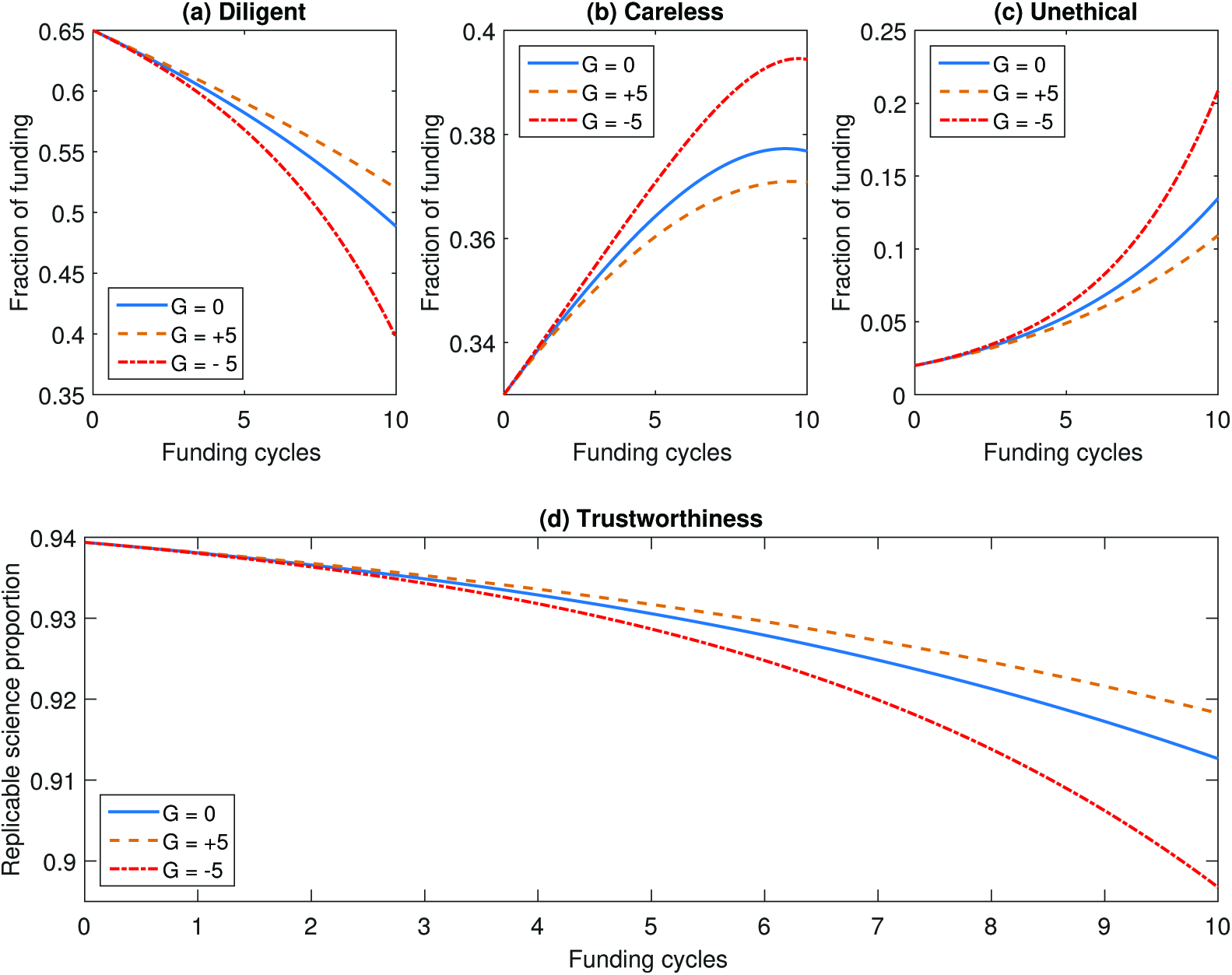
The impact of funding increases and decreases on funding allocation and science trustworthiness. Figures (a) to (c) depict the absolute proportion of funding resources allocated to diligent, careless and unethical cohorts when funding changes at rates of 0, 5 and −5 per cycle respectively.

### Impact of increased fraud detection

Figure 3 depicts the impact of aggressive fraud detection and punishment. Increased fraud detection seems to improve science trustworthiness, but *η* has to be very high in practice to have a substantial impact on the proportion of funding allocated to unethical cohorts. Negating growth, the funding allocation to this group would only be expected to decrease with time if

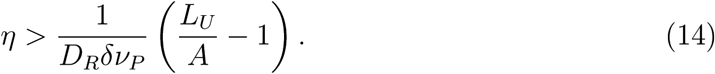

**Figure 3:**
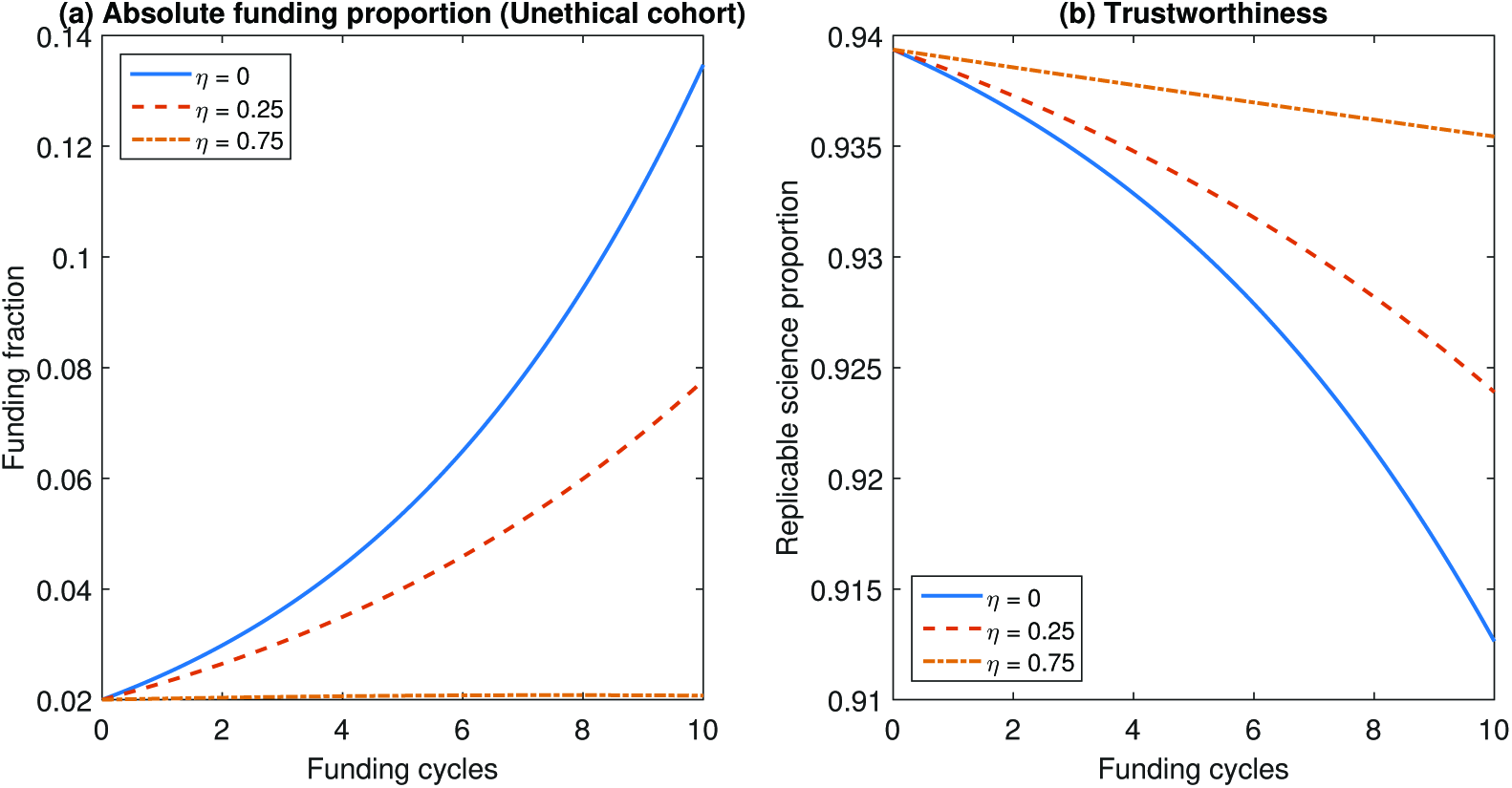
The impact of strict fraud detection / penalization. (a) Increasing the rate at which fraud is detected limits the amount of resources garnered by unethical cohorts, but (b) high values of *η* are required to markedly improve the trustworthiness of published science.

In practice this is quite high, and for the values in supplementary table 1, a value of *η* > 0.7688 would be required to fully diminish funding to this cohort in time.

### Impact of rewarding diligence

By inspect ion, it is straight-forward to s how that for the amou nt of funding held by the diligent coho rt to stay the same or increase, then the conditio n on *R_W_* is

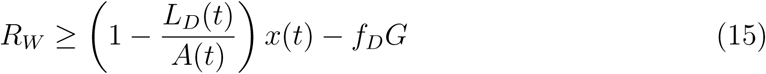

though in practice for most situations, *R_W_* will have to be much greater than this minimum value. For the example depicted in figure 4, a large reward for diligence (*R_W_* = 10) substantially increases the funds awarded to the diligent cohort. However, reproducibility still falls slowly if the unethical cohort are not removed. It is possible to both reward diligence and punish fraud, which can improve trustworthiness, illustrated in figure 4.

**Figure 4:**
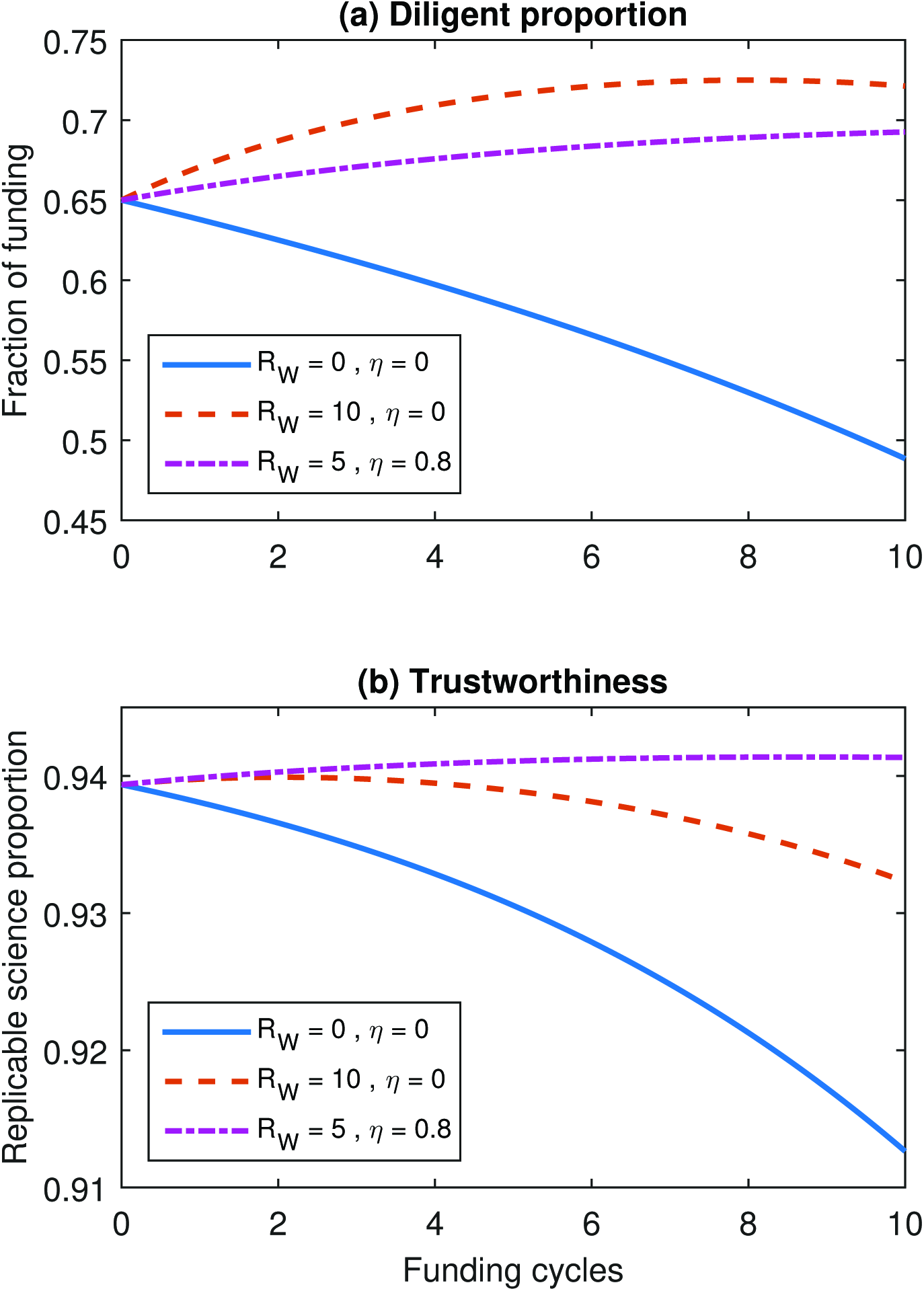
The impact of rewarding researcher for diligence. This improves the proportion of funding allocated to diligent researchers, but to improve science trustworthiness still requires non-zero values of *η* under this schema.

### Impact of the positive publication weighing

To simulate how published science fares under the rather artificial fixation of top-tier journals with positive novel results, figure 5 depicts how funding is allocated and science trustworthiness changes with varying values for *B*. In this simulation, *β_D_* = *β_C_* = *β_D_* = 0.5 when publications were *B* dependent. For *B*-independence, null results were as likely to be published so all were submitted and thus *β_D_* = *β_C_* = *β_D_* = 1. Higher values of *B* lead to perverse rewarding of false positives and fraudulent results at the expense of diligence science. Best outcome for science trustworthiness was observed when journals were simulated as completely agnostic to findings.

**Figure 5:**
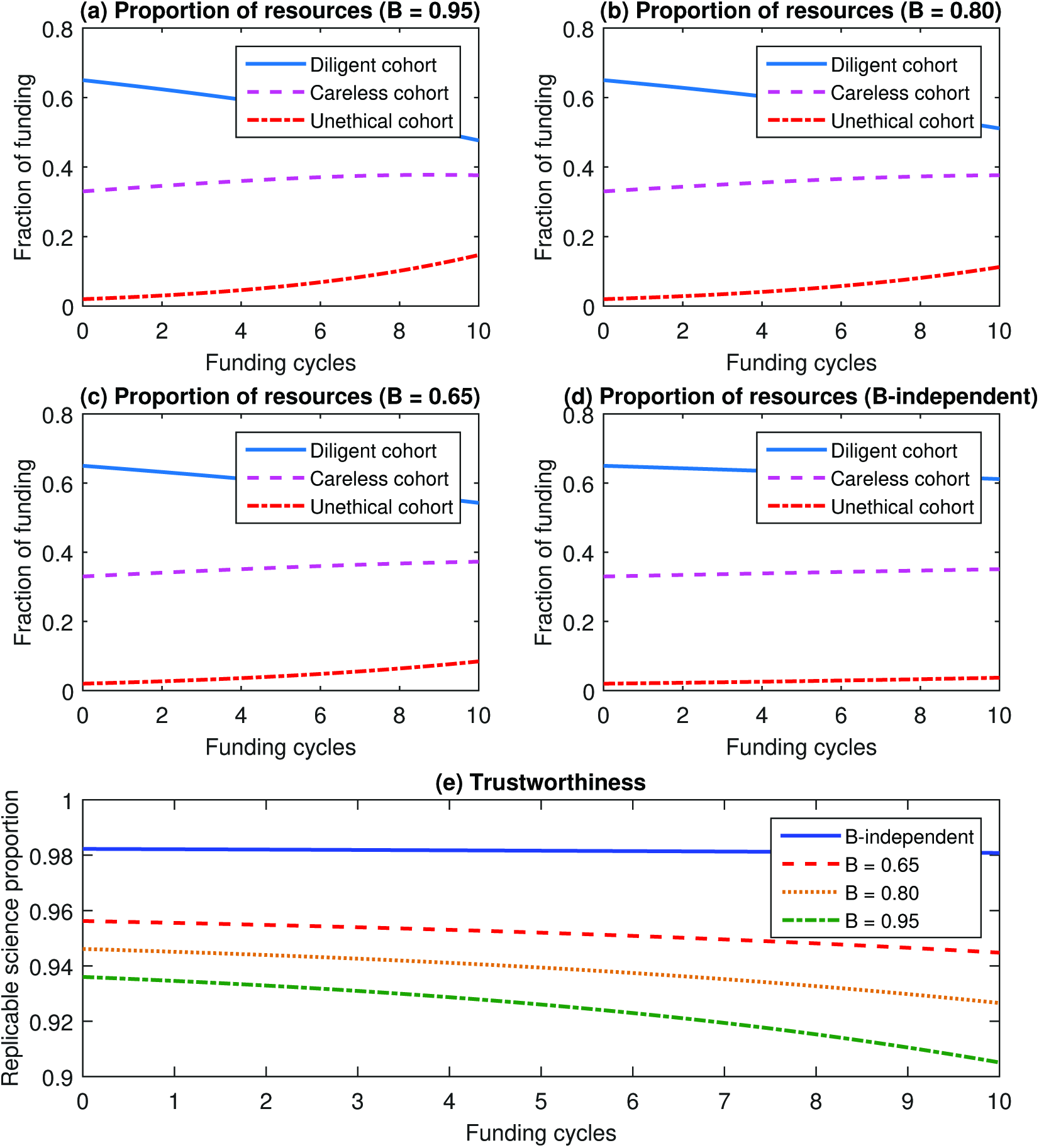
The impact of positive publication weighing on the trustworthiness of published science. (a), (b) and (c) s how funding allocation 95%, 80%, 65% of published results are positive respectively, whilst (d) depicts the situation when publications are completely agnostic. Science trustworthiness of all these scenarios in shown in figure (e), suggesting best trustworthiness obtained when journals were completely agnostic to whether a result was positive or null.

## Discussion

The model presented is a simplification of a complex eco-system, but gives some insight into what factors shape scientific trustworthiness. A full discussion of assumptions, further findings and avenues for future work are included in supplement 1. The model suggests that a fixation in top-tier journals significant or positive findings tends to drive trustworthiness of published science down, and is more likely to select for false positives and fraudulent results. In our simulations, best outcome was obtained by simply paying no heed to whether a result was significant or not. This is akin to the model used by many emerging open access peer-reviewed journals such as PLOS One, who have a policy of accepting any work provided it is scientifically rigorous. Our simulation suggests this model of publishing should improve science trustworthiness, and it is encouraging that many other publishers are taking this approach too, including Royal Society Open Science and Nature Scientific Reports. As of 2017, Scientific Reports has surpassed PLOS One as the world's biggest mega-journal [32].

While this is encouraging in one respect, there is still a perception that such journals are for ‘trivial’ or unimportant results, and that positive or important results should still go to a few journals with extreme competition for space. Empirical evaluations show that small studies published in top-impact journals have markedly exaggerated results on average compared with similar studies on the same questions published in journals of lesser impact factor [33]. This suggests that the pressure to publish in these flagship journals may still be very real, despite the option of publishing in less competitive journals. The analysis here suggests that science trustworthiness is affected too by changes in funding resources, and that when an increase of funding improves the over-all trustworthiness of science, as depicted in figure 2. Conversely when this is diminished, the increased competition on scientists appears to create conditions when false positives and dubious results are more likely to be selected for and rewarded. This is a natural consequence of the model.

One curious result persistently seen in the model was that diligent researchers are unfairly affected by careless or unethical conduct, with avoidable false positives or unethical publications garnering disproportionate reward at their expense. Simply increasing fraud detection doesn't do much to stop this, as careless researchers benefit from the gap in the market, out-producing their diligent colleagues, as seen in figure 3. This appears to be an unfortunate and seemingly unavoidable consequence of a ‘publish or perish’ system. However, in good scie ntific enviro nments carelessness would be sooner or later detected and potentially penalized. We can estimate how much of a penalty for carelessness or reward for diligence we need so as to inverse the worsening trends that we observe, by manipulating equations similar to the manner outlined for unethical conduct. However, this approach risks being ruthlessly punitive, punishing honest mistakes with the same severity reserved for the most egregious abuses of scientific trust.

While a penalty for carelessness has intuitive appeal, distinguishing between honest and careless errors is fraught with difficulty. As has been argued elsewhere [30, 31], rewarding diligence is perhaps a better way to ensure researchers do not suffer for good conduct. A simple model of this is shown in figure 4, and indeed this suggests rewarding diligence improves the proportion of funding allocated to diligent groups. However, it requires some penalty for bad conduct to keep unethical cohorts from benefiting at the expense of others. In practice this level of detection appears to have to be relatively high, which of course would require considerable resources to achieve.It should be noted too that the false positive rate of a field has a significant impact on science trustworthiness, as illustrated in figure 1. A high type II error rate provides ample cover for an unethical researcher to cheat without overt fear of detection [22, 26], perhaps explaining the elevated prevalence of dubious practice in biomedical science [22] in particular.

Future work with more sophisticated models could explore how best to implement these and other possible interventions designed to improve science trustworthiness. For instance, trustworthiness as a function of positive publication bias (B) and fraud detection rate (*η*) could be computed and optimization approaches could be applied to determine the optimal combination of B and *η* to improve science trustworthiness. These parameters can be somewhat influenced by large academic societies, government agencies, or independent foundations for instance, who could fund efforts to detect fraud in published work and support research concerning null results.

It's also important to note that the model results pivot explicitly on the assumption that scientists are forced to operate under a ‘publish-or-perish’ regime, and rewarded solely on output. Thus there is another way to improve the trustworthiness of published science - while publications are indeed one measure of productivity, they are not necessarily the sole measure. While a much harder aspect to gauge, trustworthiness is more fundamentally important. For their part, scientific journals should realize that issues such as replication and null findings are equally vital to good science as eye-catching ‘new’ results. This is slowly beginning to be recognized, with some groups coming to the forefront of championing reproducible research methods [34]. The consequences detailed in this manuscript only arise when publishing quantity is the dominant measure of an academic's worth, but in reality this should only be one consideration amongst many. The model suggests that if publishing is the sole criteria under which academics are judged, then dubious conduct can thrive.

We accordingly need to address alternative ways to assess researchers, and to encourage judicious diligence over dubious publishing. The model outlined here is far from complete, but yields some insights into the factors that shape the trustworthiness of published science. There is already evidence that pressure to publish is driving researcher burn-out and cynicism in published research [35], negatively affecting both research and the researchers themselves [36, 37]. Other studies have not found a clear association of some productivity incentives with bias [38], but these incentives may be confounded in that sometimes they coexist with other features and research practices that tend to increase also quality of research, rather than just quantity of publications. Crucially, bogus findings risk undermining public confidence in science. Amongst notable examples [39–41], the fraudulent *Lancet* MMR-Autism paper [42] is especially infamous, remaining a cornerstone of anti-vaccine narratives [43].

Publishing is not intrinsically flawed, and conversely complete, unbiased publication is essential for scientific progress. We need a better understanding of the factors driving publication and productivity-related behaviors. This is key not only to appreciating the exceptional pressures wrought upon researchers by a strict publish-or-perish imposition, but to improving science itself. This would not only benefit those working in the field, but is crucial if public trust in science is to be maintained.

## Acknowledgements

DRG would like to thank Dr. Alex G. Fletcher and Dr David Basanta for their useful feedback and thoughtful imbibing.DRG would also like to thank Prof. David Colquohoun FRS for his thoughtful discussion and advice.

